# Overview: Modeling Heterogeneous Tumor Tissue as a Multiphase Material

**DOI:** 10.1101/031534

**Authors:** Hermann B. Frieboes

## Abstract

Tumors are typically heterogeneous tissues comprised of multiple cell species in addition to extra-cellular matrix (ECM) and water fluid. It is difficult to model these components at the tissue (10^−3^–10^−2^m) scale, where individual cells cannot be represented without prohibitive computational burden. Assuming that same-kind components tend to cluster together, a multiphase approach can be applied to represent heterogeneous tumor tissue at this larger physical scale. This method enables simulating mixture of elements within tissues, e.g., geno-/phenotypic heterogeneity underlying mutation- or microenvironment-driven tumor progression. Further, by not explicitly tracking interfaces, this methodology facilitates realistic modeling of tissue in 3-D.

## I. Representation of tumor tissue heterogeneity

With a multiphase approach, a solid tumor is modeled as a saturated medium consisting of at least one solid phase (cells, ECM, etc.) and one liquid phase (fluid) [1]. The approach can be generalized to represent additional phases to describe multiple tissue elements. The volume fractions or mass at a given location describe the relative amounts of the different elements. The system is governed by mass and momentum balance equations for each phase, interphase mass and momentum exchange, along with appropriate constitutive laws to close the equations [1]. The approach does not require enforcing complicated boundary conditions between elements. This method stands in contrast to modeling using *sharp boundaries*, in which tumor elements at the tissue scale are delineated from each other without mixing, hence potentially representing a challenge to properly simulate the tumor biological complexity.

Many multiphase mixture models have been developed to account for cell type and mechanical response heterogeneity of the solid and liquid tumor phases. Work includes [1–39] in the past decade and associated earlier-date references.

## II. Illustrative Results of application of Method

Multiphase tumor models in 3-D [39] have been combined with models of tumor-induced angiogenesis [40–41] to fully couple heterogeneous tumor growth with vascularization [37–38, 42–44]. The nonlinear effects of cell-to-cell adhesion as well as taxis-inducing chemical/molecular species are implemented by using an approach based on energy variation [1]. This means that the energy within the tumor system accounts for all the processes to be modeled. In particular, adhesion is represented through an interaction energy that leads to narrow transition layers originating from differential adhesive forces among the cell species [1]. The resulting diffuse-interface mixture equations for the cell-species volume fractions are a coupled system which includes fourth-order nonlinear advection–reaction–diffusion equations of Cahn–Hilliard-type [45]. Cell substrate elements (e.g., oxygen) are included via reaction–diffusion equations. This framework allows for a detailed description of solid tumor growth without the need to track sharp boundaries between the tumor constituent elements. It also firmly links the dependence of cell–cell and cell–matrix adhesion on cell geno- and phenotype and on local microenvironmental conditions (such as levels of oxygen) [1].

### A. Simulation of vascularized tumor growth

Fig. 1(left) simulates growth of a vascularized tumor [42]. Hypoxia leads to necrosis in the interior (darker color), with tumor angiogenic factors (TAF) diffusing from the interior to the surroundings to stimulate new capillaries (small lines) from pre-existing vasculature (not shown). Endothelial cells proliferate up the TAF gradient, forming branches and then loops to conduct blood (darker lines). Vessels are uniformly distributed around and inside the lesion. In time, the tumor becomes asymmetric, influenced by heterogeneity in cell proliferation and death, which is in turn based on availability of cell substrates in the microenvironment as a function of the vasculature. Fig. 1(right) shows that viable tissue cuffs around the vessels, as observed experimentally [46] and clinically [37–38], with areas distal from conducting vessels undergoing necrosis.

**Figure 1.**
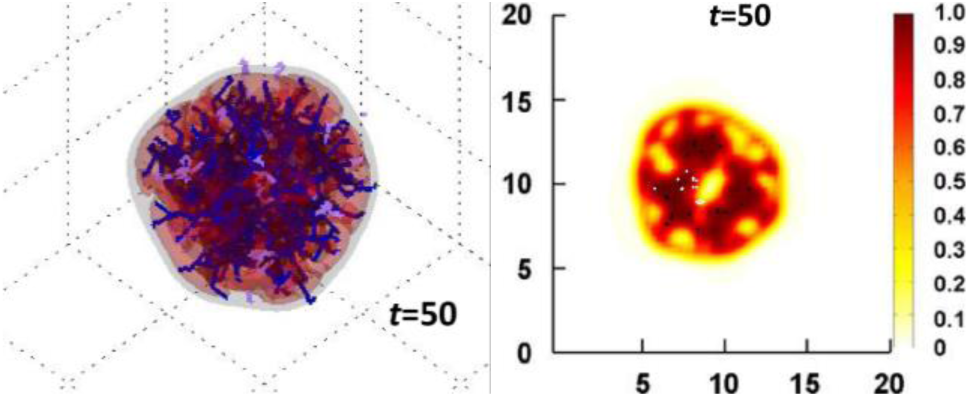
Simulation of vascularized tumor with one viable cell species [42]. Left: Viable tumor tissue (orange/ red) is shown in 3-D contours representing density values of 0.1, 0.2, and 0.6 (min.:0.0; max.:1.0). Conducting vessels: blue; non-conducting: gray. Time unit = 1 day; grid length = 200μm. Right: Slice of tumor (plane x=10) shows that viable tissue cuffs around the vessel locations (darker areas). Color coding: density of viable tissue; highest=1.0; unit length = 100μm.

### B. Simulation of multiple viable tumor species

Fig. 2 shows a tumor that originally started as a small, round avascular tumor nodule with one viable species [42]. As in the previous example, hypoxia leads to necrosis in the interior, with TAF diffusing from the interior to the surroundings to stimulate new capillaries that eventually conduct blood (darker lines). A mutated species (red) with equal oxygen uptake as the original species (gray) is made to appear randomly, expressing a phenotype which upregulates migration when subjected to hypoxia, as would occur in regions distal from conducting vessels. This species down-regulates proliferation until reaching regions with sufficient oxygen, such as the tumor/host interface. Over time, the migratory phenotype by itself does not seem to provide an overriding advantage to the mutated species. Since migration is down-regulated once adequate oxygenation is reached, this example illustrates that even if migration enhances species survival, it may lower net proliferation for the tumor as a whole.

**Figure 2.**
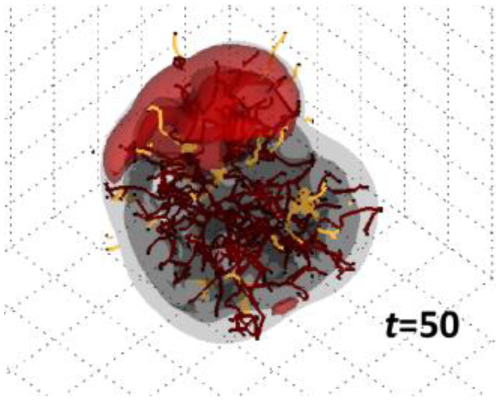
Simulation of two viable tumor species, one with a migratory phenotype when hypoxic [42]. The two species compete for oxygen in the microenvironment, with the migratory species (red) aggregating on one side. Viable tumor tissue (gray) is shown in 3D contours representing density values of 0.1, 0.2, and 0.6 (min.:0.0; max.:1.0). Conducting vessels: brown; non-conducting: yellow. Time unit = 1 day; grid length = 200μm.

## III. Quick Guide to the Methods

We summarize the general formulation stated in [1,39,42].

### A. Conservation of Mass

The dependent variables in a model with (*N*+1) species are:

○ Volume fractions of the species *ϕ*_0_,…, *ϕ_N_*
○ Densities of the species *ρ*_0_*,…, ρ_N_*
○ Extra-cellular (solid) pressure *p*, and
○ Component velocities **u**_0_,…,**u**_N_

It is assumed that there are no voids and therefore, 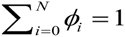, and that the densities are constant. Without loss of generality, *i*=0 denotes the water fluid component, *i*=1,…,*N*-1 denote the tumor species components, and *i=N* denotes the host species component. Assuming the water volume fraction is constant in time and space, the volume fraction of the solid components is also constant, 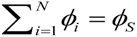.

The volume fractions are assumed continuous in a domain Ω, containing both tumor and host domains. The fractions satisfy the mass conservation advection-reaction-diffusion equations:

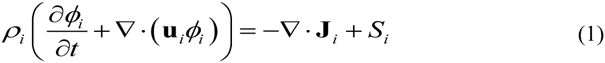

where J*_i_* are fluxes accounting for the mechanical interactions between the species. The source terms *S_i_* account for the mass exchange between components and mass gain and loss due to cell proliferation and lysing, respectively. The mass of the mixture is conserved only if

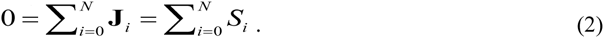

An approach from continuum thermodynamics (e.g., [47–48]) introduces the Helmholtz free energy of component interactions (e.g., adhesion). For the *i_th_* component:

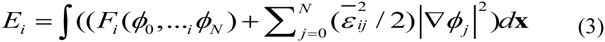

where the first term models bulk energy due to local interactions while the second term represents longer range interactions. 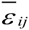 are non-negative constants such that 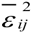 has units of energy per unit length. The total energy is 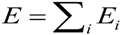.

If the volume fraction of water is assumed constant in space and time, then the volume fraction of the solid components is also constant. Thermodynamically consistent fluxes may then be taken as in the generalized Fick’s Law:

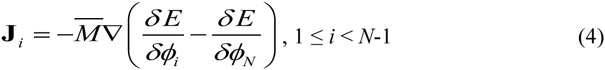

and 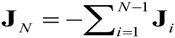. The mobility 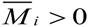, while *δE* / *δϕ_i_* are variational derivatives of the total energy E:

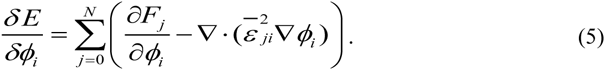

The resulting generalized Darcy laws for the component velocities are:

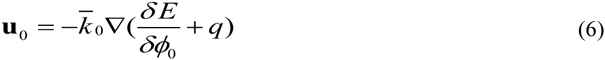

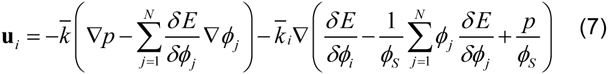

where *q* is water pressure and *p* is solid pressure. The terms dependent on *δE* / *δϕ_j_* represent the excess force due to adhesion arising from cell-cell and cell-host interactions. The coefficients 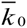, 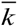, and 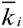 are positive definite motility matrices reflecting the response of the water and cell species to pressure gradients. The motilities encapsulate the combined effect of cell-cell and cell-matrix adhesion. Equations (4), (6), and (7) are constitutive laws that guarantee that in the absence of mass sources, the adhesion energy is non-increasing in time as the fields evolve.

### B. Type of settings in which this method are useful

This method has been used to model avascular as well as vascularized tumor growth, but in general it can also be applied to study biological problems involving different components interacting and intermixing with each other, such as occurs during embryogenesis and organ development. The method is especially useful to simulate tissue-scale interactions in 3-D without having to track individual cells or molecular components, and the interfaces between them.

